# Neuroligin affects metastatic performance of CD66+ cells: Theoretical analysis of Multi-tissue RNA expressions through System modeling based Network mapping under *in-vitro* constraints

**DOI:** 10.1101/2020.07.29.226274

**Authors:** Om Prakash

## Abstract

Present study deals with theoretical analysis of multi-tissue RNA expressions through system model based network mapping under the constraints of *in-vitro* experiments. Here, effect of Neuroligin was evaluated on the performance of metastatic behavior of CD66+ cells of human female reproductive cell lines. Total 10 genes and 14 cell lines were considered in the study. Clustering of cell lines was done on the basis of network mapping under the same homeostatic constraints. Neuroligin was found to affect metastatic performance of CD66+ cells. Two CD66+ cell lines hTERT-HME1 and SK-BR-3 were found to most effective, in similar manner as brain cell lines as U-138MG & U-87MG.

## INTRODUCTION

Biology of malignancy of cancer cells are directly related with cell adhesion molecules and transition of clustered epithelial cells into free-to-move mesenchymal cells. Two main genes NLGN4X (for neuronal adhesion) (***Henderson et. al., 2017***) and CD66 (for cervical cancers) (***Ammothumkandy et. al., 2016; Arabzadeh et. al. 2016; Oliveira et. al. 2018; Wicklein et. al., 2018; Calinescu et. al. 2018; Wang et. al. 2019***) are known for their effect on malignancy in epithelial cancer cells (***Wegwitz et. al. 2016; Zhuo et. al. 2016***). In some studies, impact of NLGN4X was also observed on lymphoid tissue. Epithelial to mesenchymal transitions are related with prolyl 4-hydroxylase 9 (P4H9) and glycosylation at molecular level (***Yokoyama et. al., 2015***).

Cellular behavior depends on differential gene expression. Since each of the gene expressions is differentially affected by the co-expressed genes. Interdependency of differentially expressed genes is the key motivation behind the understanding of systems biology. Components of system behave towards each other by some naturally defined rules, which are not completely understood. These rules define the blueprint for cellular homeostasis mechanism. This Property protects cell survival by reorganizing strength of gene expression even in the acidic environment as well as in hypoxic conditions. This cellular homeostasis also support for the survival of cell during cellular transition as well as migration. Therefore information for tissue level expressions of genes can be used for understanding the behavior of cells for metastasis. One of the most prominent examples is Neuroligin 4X (NLGN4X)(GSE96632). It is a postsynaptic cell addition molecule and play role in various neuronal disorders. Its high expression in non-metastatic melanomas and low expression in metastatic melanomas has been observed. Knockdown of NLGN4X was found to cause expressional changes in HIF1 pathway. Metastatic potential of melanomas were also found to be linked with their ability to recruit lymphatic vessels. Therefore it is assumed that neuroligin may affect the metastatic behavior of various cell lines.

Therefore to understand metastatic behavior in cancer cells, differential gene expressions were observed in various cell lines from normal, brain and CD66+ female reproductive system. Here impact of neuroligin was observed on metastatic behavior of CD66+ female reproductive cell lines.

## MATERIALS & METHODS

### Dataset

Initial data was collected as RNA expressions for 10 genes observed in 14 cell lines including: HaCaT (normal); Brain cell lines AF22, SH-SY5Y, U-138MG, U-251MG, U-87MG; and female reproductive organ cell lines AN3-CA, EFO-21, HeLa, hTERT-HME1, MCF7, SiHa, SK-BR-3, T-47d. Genes considered for study were: Neuroligin (NLGN4X); CD66 (CEACAM8); homeostasis observing control genes: GAPDH, ACTB; NLGN4X influenced genes involved in metastasis : HIF1A, ATP10A; and CD66 influenced genes involved in metastasis HAS1, MFHAS1, DDX18, CSN1S1. RNA expressions of genes for respective cell lines have been tabulated in **Table 1**.

**Table 1.**
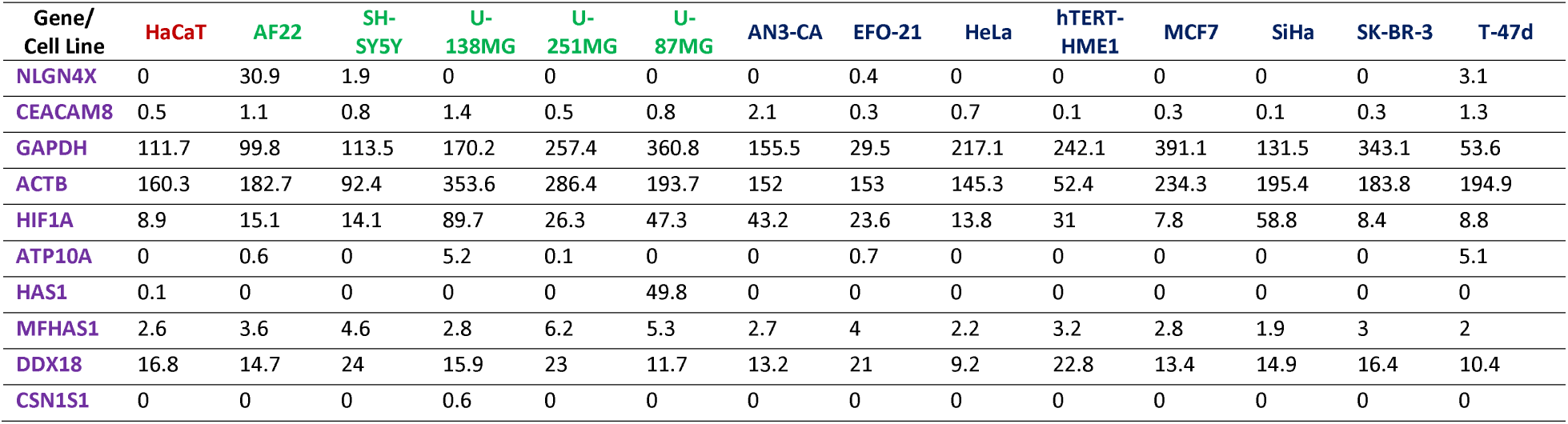
RNA expressions of genes for respective cell lines (*Source: Human Protein Atlas*)

### System model & Network mapping

Next Generation Sequencing provides details about the genes being expressed as well as differential gene expression. The data shows temporal expression. Various extends of temporal expressions have been recorded in various databases as Human Protein Atlas. It has collection of multiple cell lines as well as respective genes. In present study a set of multiple gene RNA expression values from various cell lines were used to create a systems model. This systems model was motivated for understanding the metastatic behavior of various CD66+ cell lines under the influence of neuroligin expression. In this study the framework of homeostasis maintenance was used as a reference point to evaluate performance of various cell lines. 03 categories of cell lines (normal, neuronal and female system cell lines) were used in systems modeling. Furthermore these cell lines were compared on the basis of 10 different gene expressions as defined.

***For systems modeling***, normalized temporal expressions of RNA was processed with Gamma distribution function, which was further utilized for development of multi-gene system model. This model was simulated for all combinations of gene pairs for differential expression. This simulation was adopted in bidirectional rate combinations between each pair of gene expressions. Further absolute difference between expressions of two directions was observed on the ground of time movement. The modeling process was performed with in-house Python programs.

***For network graph***, correlation coefficient between each gene-pair was used as edge-weight. System’s gene network graphs were drawn for each cell line separately. This set of cell lines were further clustered on the ground of maintenance of homeostasis in relation of unit correlation with metastasis representing genes from neuronal as well as female reproductive system cell lines. The modeling process was performed with in-house Python programs with Networkx package.

## RESULTS & DISCUSSION

Initial study framework was defined with 10 genes and 14 cell lines. Study has been performed to cluster the metastatic behavior of cell lines under the influence of expression of neuroligin. This study was performed theoretically but constraints of experimental limitations as well as biological rules were adopted from *in-vitro* experiments.

As the multiple studies showed that gene expressions follow Gamma distribution, therefore in this study RNA expression was simulated according to it. Expression of each gene was calculated in two directions of highest and lowest expressions. Gene expression was populated with time including expression limitations identified in experimental studies. Gene interaction was defined by absolute correlation difference established between bi-directional expressions of gene-pairs simulated on time unit of 1 to 10. All combinations of gene interactions were plotted as network graph. Separate network graph was plotted for each cell line. A homeostasis criterion was defined by the relation of GAPDH and ACTB with NLGN4X and CD66. All combinations of gene interaction in network graph were filtered on the basis of homeostasis criteria. Per unit homeostasis in combination of NLGN4X and CD66 was calculated; and cell lines where clustered on the basis of minimum find maximum value of per unit homeostasis established.

This set of cell lines where further clustered on the ground of maintenance of homeostasis in relation of unit correlation with metastasis representing jeans from neuronal as well as female reproductive system cell lines. Ultimately, Neuroligin found to affect metastatic performance of CD66+ cells. Two CD66+ cell lines hTERT-HME1 and SK-BR-3 were found to most effectively behave in similar manner as brain cell lines as U-138MG & U-87MG, under the same homeostatic constraints. Other cell lines as EFO-21, MCF7, SiHa & T-47d were found to be less affective; while AN3-CA did not showed any effect from NLGN4X (**Figure 1 & 2**). Gene interaction networks and pattern gene expressions have been plotted in **Table 2 & 3**.

**Table 2.**
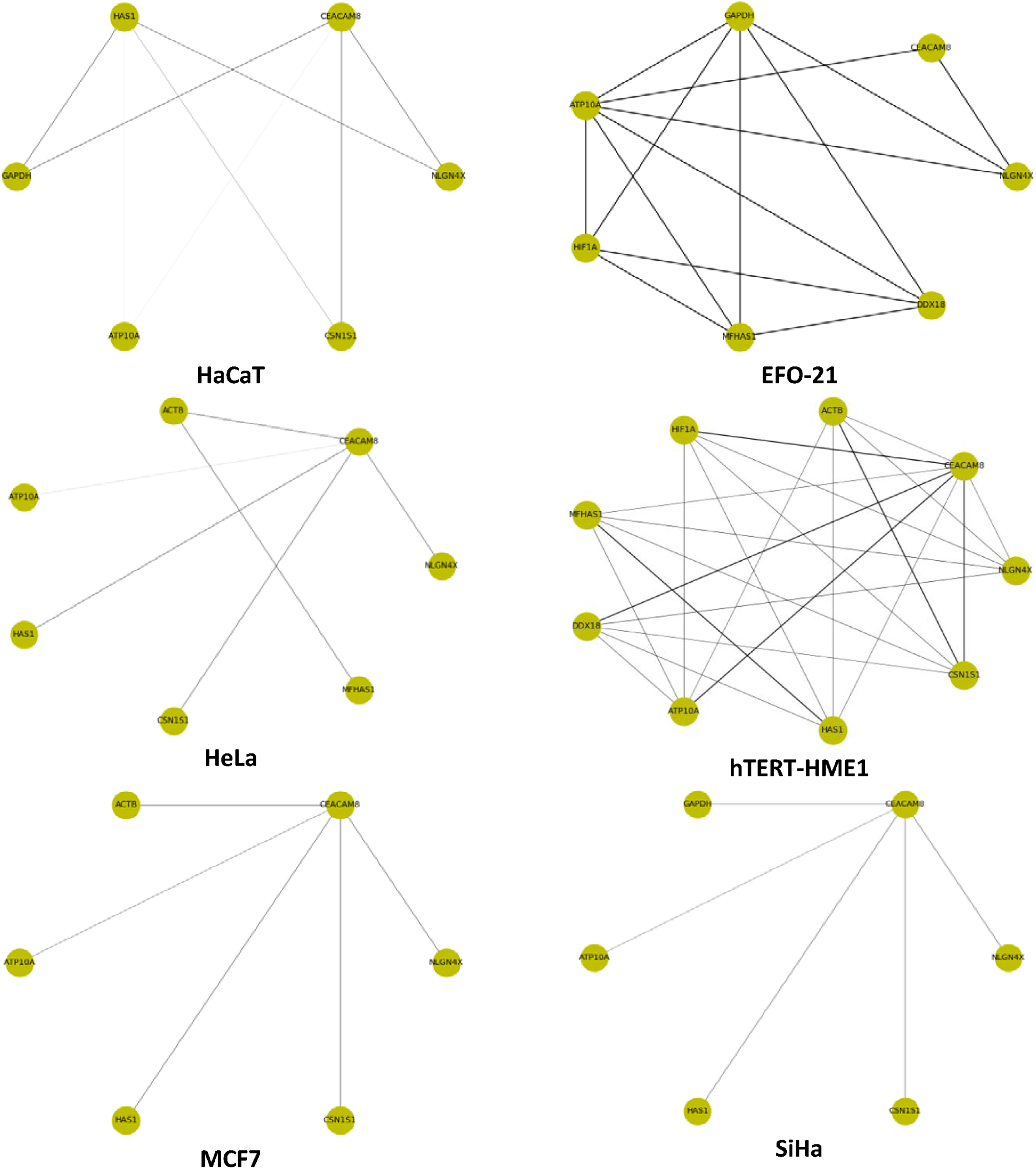
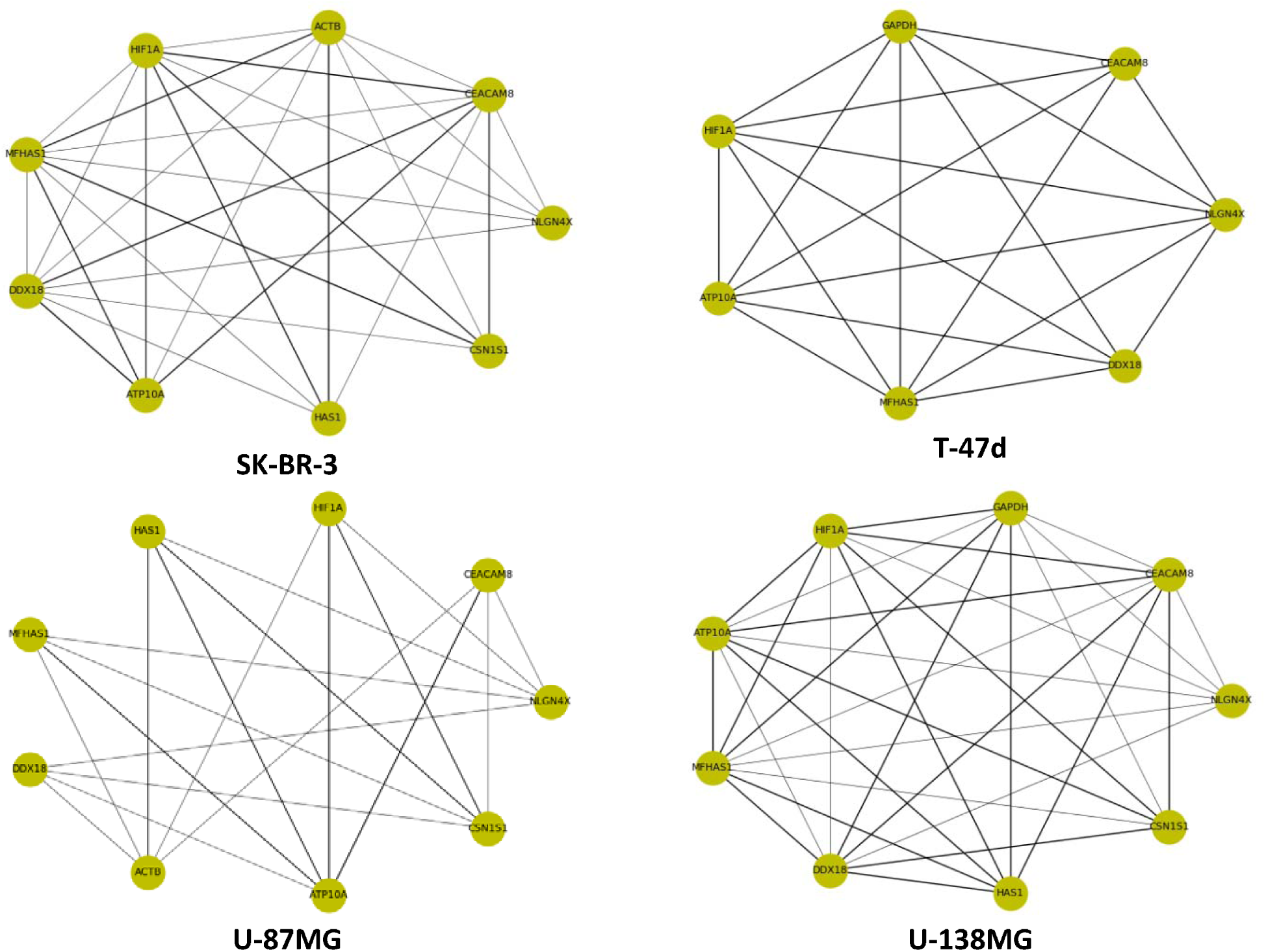
Cell line wise filtered networks from systems models

**Table 3.**
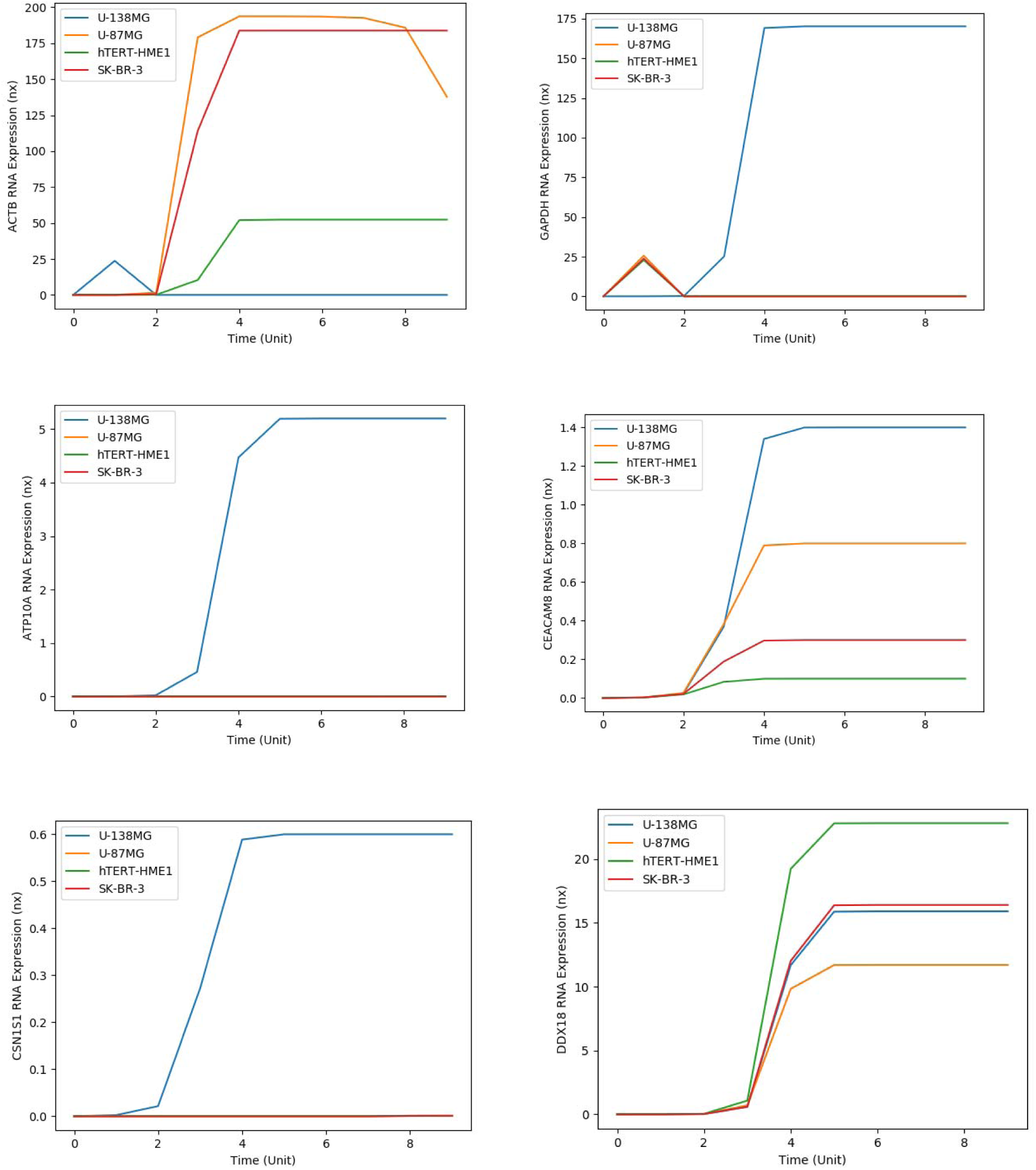
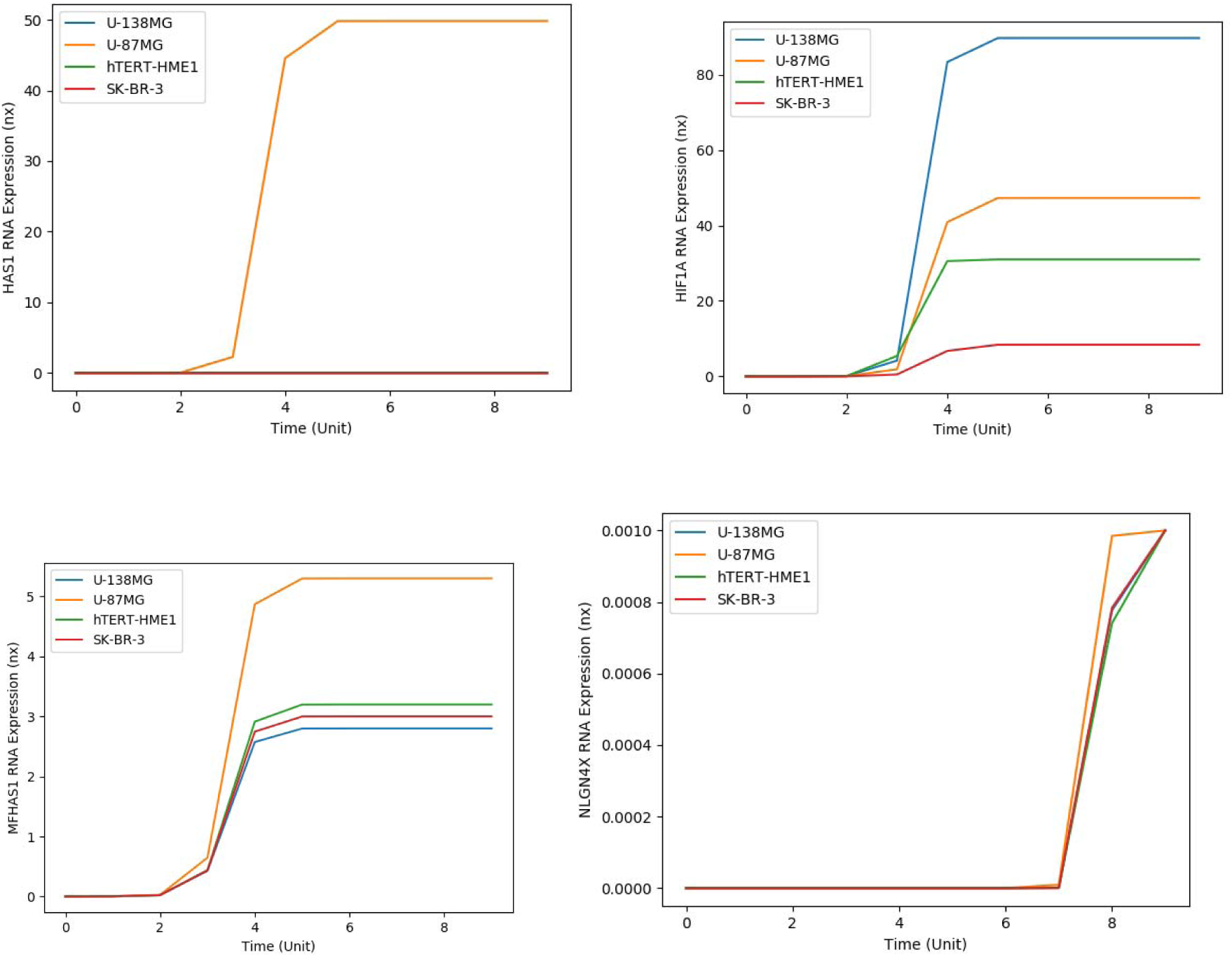
Simulated gene expressions from various cell lines filtered networks from systems models

**Figure 1.**
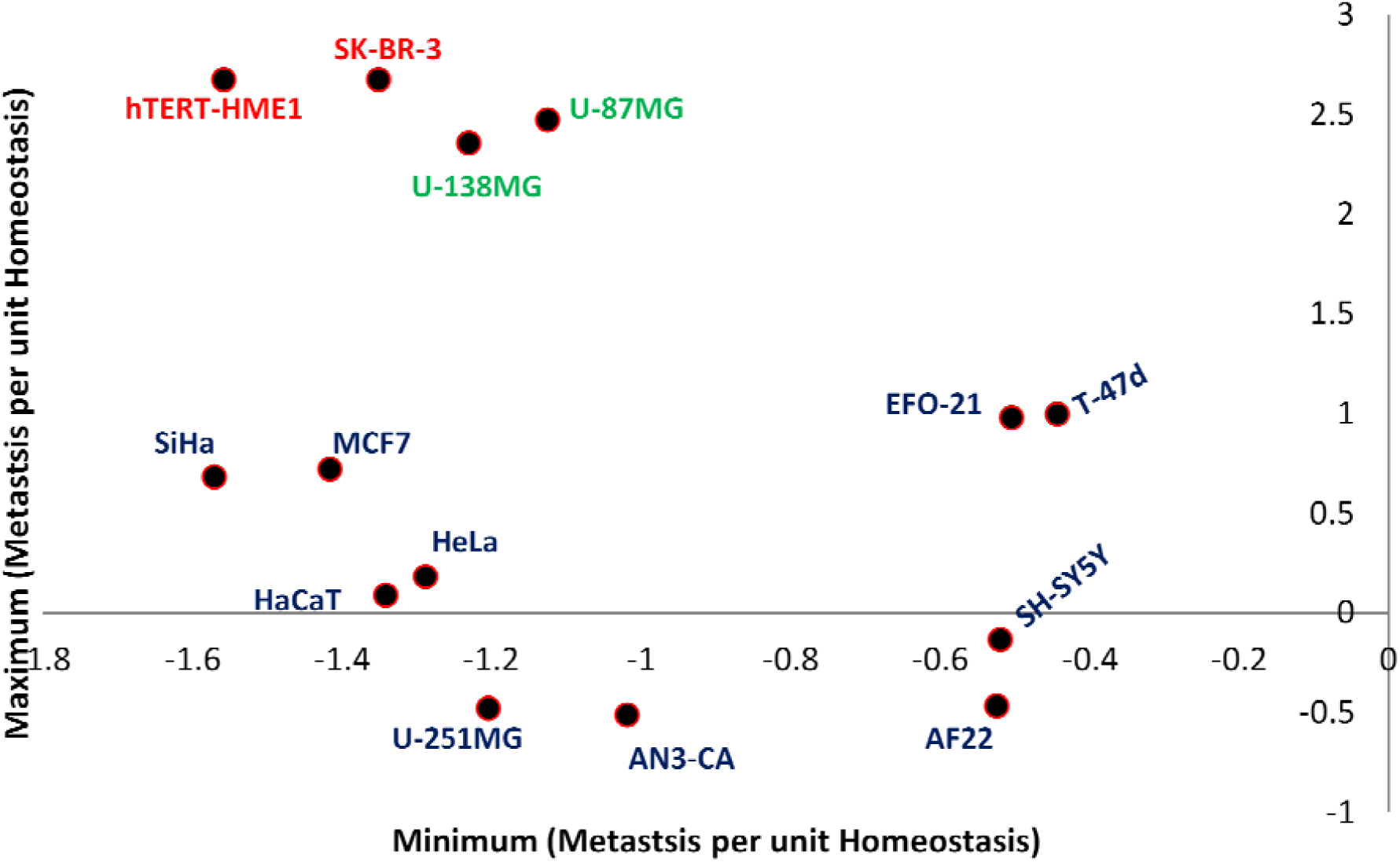
Clustered cell lines which are most influenced with NLGN4X under homeostasis constraints. Two CD66+ cell lines hTERT-HME1 and SK-BR-3 were found to most effectively behave in similar manner as brain cell lines as U-138MG & U-87MG.

**Figure 2.**
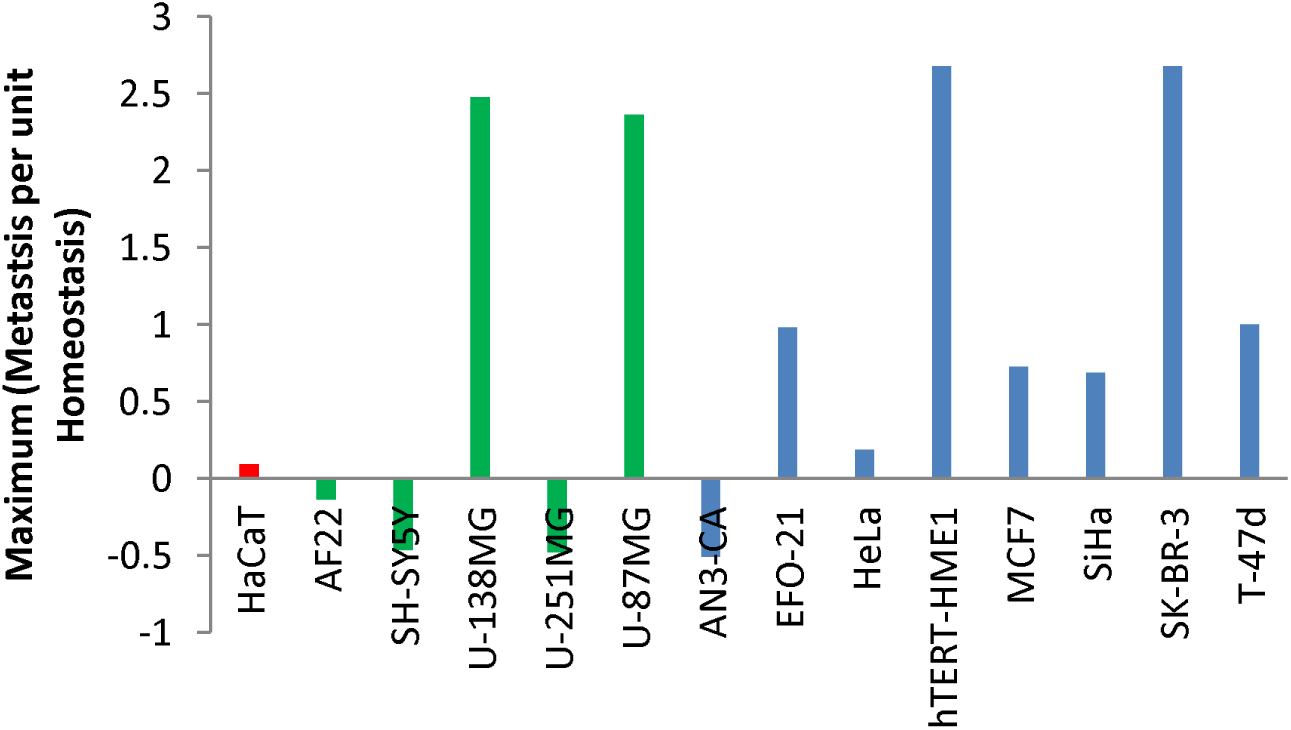
Clustered brain (Green) and female reproductive cell lines (Blue) along with HaCaT (Red). These cell lines bars are showing effect influenced with NLGN4X under homeostasis constraints.

## CONCLUSION

Neuroligin found to affect metastatic performance of CD66+ cells. Two CD66+ cell lines hTERT-HME1 and SK-BR-3 were found to most effectively behave in similar manner as brain cell lines as U-138MG & U-87MG, under the same homeostatic constraints. Other cell lines as EFO-21, MCF7, SiHa & T-47d were found to be less affective; while AN3-CA did not showed any effect from NLGN4X.

## ACKNOWLEDGEMENT

Author express gratitude to *The Institute of Mathematical Sciences*, Chennai-600113, India for providing research facilities as well as DAE Post-Doctoral Fellowship (PDF 214).

